# ERGA-BGE reference genomes of *Hyalomma lusitanicum* and its obligate *Francisella* endosymbiont as a genomic resource for One Health research

**DOI:** 10.64898/2026.05.27.728183

**Authors:** Juan E. Uribe, Juan S. Echeverry-Pérez, Félix Valcárcel, A. Sonia Olmeda, María Sánchez Sánchez, José M. Tercero, Nuria Escudero, Rosa Fernández, Astrid Böhne, Rita Monteiro, Marta Gut, Laura Aguilera, Francisco Câmara Ferreira, Fernando Cruz, Jèssica Gómez-Garrido, Tyler S. Alioto, Christian de Guttry

## Abstract

*Hyalomma lusitanicum* is a characteristic tick species of the western Mediterranean region, with a well-established distribution across the Iberian Peninsula. It is strongly associated with wild ungulates, particularly red deer, as well as livestock, to which it can transmit a wide range of pathogens, including viruses, bacteria, and protozoa. Here, we present three genomic resources for *H. lusitanicum*: a scaffold-scale nuclear genome, the complete mitochondrial genome, and the complete genome of its associated *Francisella* bacterial endosymbiont. The nuclear genome assembly spans 1.81 Gb and comprises 59 scaffolds, with a scaffold N50 of 153.6 Mb (L50 = 5) and no gaps, indicating high contiguity and completeness with a gene annotation completeness BUSCO score of 97.1 %. Genome annotation of the nuclear assembly identified 20,638 protein-coding genes, 1,422 non-coding genes, and 5,775 pseudogenes. A total of 18 scaffolds were assembled as putative chromosomes, exceeding the 11 chromosomes inferred as ancestral; however, synteny analyses suggest that several scaffolds likely represent fragmented portions of the same chromosome, probably due to incomplete Hi-C scaffolding. Despite this, the assembly represents one of the most complete tick nuclear genomes generated to date. In addition, we report the complete genome of a *Francisella* endosymbiont (1.51 Mb, 1,679 genes), characterized by a high proportion of pseudogenes and reduced genome size, consistent with patterns of genome reduction associated with obligate symbiosis. Together, these genomic resources provide a framework to investigate local adaptation and host-symbiont evolution, and to support improved surveillance, control, and management strategies for species of public health relevance.

## INTRODUCTION

Ticks (Acari: Ixodida) are obligate hematophagous ectoparasites that require different vertebrate hosts to complete their life cycle. This close and prolonged interaction with vertebrate hosts places ticks among the most important arthropod vectors of a wide range of pathogens, including viruses, bacteria, and protozoa, causing diseases such as rickettsiosis, Lyme disease, relapsing fever borreliosis, and babesiosis, with major impacts on human and animal health worldwide (Jongejan and Uilenberg, 2024; Magnarelli, 2009). Their ecological success is tightly linked to climatic conditions, host availability, and landscape features that are currently undergoing rapid transformation under global change (Estrada-Peña et al., 2014; Ogden et al., 2016; Gortazar et al., 2014). Consequently, tick distributions are shifting north, with several species expanding into new geographic areas, increasing the risk of emerging and re-emerging tick-borne diseases (Bah et al., 2022; Da Re et al., 2025; Gandy et al., 2023; Hornok et al., 2020). In addition, the success of ticks is strongly influenced by their association with obligate and facultative endosymbionts, which can enhance host fitness by providing essential nutrients, modulating reproduction, and potentially affecting vector competence, thereby playing a key role in tick adaptation and epidemiological dynamics (Duron et al., 2018; Weiss and Aksoy, 2011).

Overall, among hard ticks (Ixodidae), the genus *Hyalomma* is of particular relevance due to its ecological plasticity, active questing behavior, and capacity to transmit high-impact pathogens such as the Crimean-Congo hemorrhagic fever virus (CCHFV), which has a 40% mortality rate –[World Health Organization (WHO)](https://www.who.int/news-room/fact-sheets/detail/crimean-congo-haemorrhagic-fever)–. The genus *Hyalomma* comprises 27 described species that are predominantly associated with arid and semi-arid environments across Africa, the Mediterranean Basin, and parts of Asia, where temperature and humidity regimes strongly influence their life cycles and seasonal activity patterns (Estrada-Peña et al., 2011; Sands et al., 2017).

*Hyalomma lusitanicum* represents one of the most characteristic *Hyalomma* species in the westernmost Mediterranean region. It exhibits a well-established distribution across the Iberian Peninsula, particularly in Mediterranean ecosystems of Spain and Portugal, and extends into adjacent localities shaped by similar climatic and ecological conditions. The species is strongly associated with wild ungulates, especially red deer, as well as livestock, facilitating its persistence in both natural and human-modified landscapes (Valcárcel et al., 2023). In the central-western Iberian Peninsula, *H. lusitanicum* occurs in sympatry with its congeneric species, *Hyalomma marginatum*, which is widely recognized as the primary vector of CCHFV in Europe (Celina et al, 2023). In Spain, nearly all reported autochthonous cases of CCHF, including six fatal cases to date, have been geographically linked to areas where both species converge (Cano et al., 2022; Baz-Flores et al., 2024). However, although *H. marginatum* is a confirmed CCHFV vector across its distribution range, viral detection in Spanish ticks has not been fully confirmed (Negredo et al., 2019). By contrast, *H. lusitanicum* is not formally recognized as a competent vector of CCHFV. Yet, its high abundance, broad host range, and ecological dominance in several endemic areas suggest that it may contribute to local virus maintenance and circulation (Negredo et al., 2019; Portillo et al., 2021). This apparent discrepancy highlights the need to better understand the ecological and evolutionary dynamics of sympatric *Hyalomma* species within multi-vector systems.

Here, we present a high-quality nuclear genome assembly of *Hyalomma lusitanicum*, which was coordinated by the European Reference Genome Atlas (ERGA) initiative’s Biodiversity Genomics Europe (BGE) project, supporting ERGA’s aims of promoting transnational cooperation to promote advances in the application of genomics technologies to protect and restore biodiversity (Mazzoni et al., 2023). In addition, we describe the complete genome of the obligate *Francisella* endosymbiont, a key bacterial partner that contributes to tick fitness by supplying essential nutrients and potentially influencing vector competence and pathogen transmission. Together, these genomic resources establish a functional framework for investigating local adaptation and for understanding the evolutionary processes underlying the epidemiology of Crimean-Congo hemorrhagic fever virus (CCHFV).

## METHODOLOGY

### Sample and sampling information

Adult specimens of *Hyalomma lusitanicum* (n=14, 7 males and 7 females) were collected from vegetation in a mesomediterranean area of central Spain (38.464297-4.45332, Fuencaliente, Ciudad Real). The ticks were collected using the dragging technique (Barandika et al., 2014). They were identified and sexed based on morphological characteristics following standard taxonomic keys for *Hyalomma*, taking into account its sexual dimorphism (Walker et al., 2003; Apanaskevich et al., 2008; Estrada-Peña et al., 2017). The biological material collected in Spain, and used to generate digital sequences, was retrieved from wildlife taxa regulated by the Spanish Royal Decree 124/2017 (https://www.boe.es/eli/rd/2017/24/124). After identification, the ticks were preserved in laboratory glass tubes with a folded paper strip, sealed with water-repellent cotton, and then placed in biosafety containers for shipment. Two males and one female were processed as follows to obtain the genome of this species.

### Vouchering information

Proxy physical reference material has been deposited in Museo Nacional de Ciencias Naturales (MNCN) https://www.mncn.csic.es under accession MNCN20.02/22139. Frozen proxy reference tissue material is also available at the same institution under accession numbers MNCN-ADN-151978 and MNCN-ADN-151979.

### Genomic processing

Previous genetic information estimated 2.79 Gb of genome size for *Hyalomma lusitanicum* based on ancestral state reconstruction, including species from other hard-tick genera with chromosome-level genome assemblies available, such as *Dermacentor, Rhipicephalus, Amblyomma*, and *Ixodes* (GoaT). The estimate is for a diploid genome with a haploid number of 10 autosomes and 1 sex chromosome (2n=22) (García et al., 2002; Tidwell et al., 2024). All information for this species was retrieved from Genomes on a Tree (Challis et al., 2023).

#### DNA/RNA extraction

DNA was extracted from two whole organisms (female and male) using the Blood & Cell Culture DNA Mini Kit (Qiagen) following the manufacturer’s instructions. DNA quantification was performed using a Qubit dsDNA BR Assay Kit (Thermo Fisher Scientific), and DNA integrity was assessed using a Fragment Analyser System (Agilent). The DNA was stored at +4 °C until sequenced.

RNA was extracted from a whole organism (SAMEA114541064) using an RNeasy Mini Kit (Qiagen) according to the manufacturer’s instructions. RNA quantification was performed using the Qubit RNA BR kit, and RNA integrity was assessed using a Bioanalyzer 2100 system (Agilent) RNA 6000 Nano Kit (Agilent). The RNA was stored at −80 °C until sequenced.

#### Libraries preparation

For long-read whole-genome sequencing, one library was prepared using the SQK-LSK114 Kit (Oxford Nanopore Technologies, ONT) (Biosample SAMEA114541056), and then sequenced across four R10.4.1 flow cells on a PromethION 24 A Series instrument (ONT) (Biosample SAMEA114541046). A short-read whole-genome sequencing library was prepared using the KAPA Hyper Prep Kit (Roche) (SAMEA114541046). The RNA library was prepared using the KAPA mRNA Hyper prep kit (Roche). All short-read libraries were sequenced on a NovaSeq 6000 instrument (2×151bp, Illumina).

### Nuclear genome assembly

The genome was assembled using the CNAG CLAWS pipeline v2.2 (Gomez-Garrido, 2024). Briefly, ONT reads were pre-processed for quality and length using Filtlong v0.2.1 (https://github.com/rrwick/Filtlong) with the options --min_length 1000 --min_mean_q 90 --target_bases 140000000000, resulting in 140 Gb of reads with an N50 of 36.4 kb. Contigs were assembled using Hifiasm v0.24.0 (Cheng et al., 2021) with the --ont option. The haplotype 2 assembly was chosen as the final assembly as it had the highest QV, the lowest level of false duplications and similar contiguity to the primary assembly. The haplotype 1 assembly was incomplete and more fragmented. BUSCO v5.4 (Manni et al., 2021) using the arachnida_odb10 lineage, and Merqury (Rhie et al., 2020) were used to estimate completeness and quality.

### Mitochondria and endosymbiont genome assembly

Long-read (ERR13381541) and short-read (ERR13765978) data from the same individual (SAMEA114541046) were quality-trimmed using fastlong and fastp, respectively (Chen, 2023; 2025). The processed reads were then mapped against the assembled and contamination-filtered nuclear genome of *Hyalomma lusitanicum* using Minimap2 (Li, 2018).

Unmapped long and short reads were extracted with samtools (Danecek et al., 2021) and subsequently assembled *de novo* using SPAdes (Prjibelski et al., 2020) with the following command: spades.py −1 ERR13765978_unmapped_R1.fq.gz −2 ERR13765978_unmapped_R2.fq.gz --pacbio ERR13381541_unmapped.fq.gz -o de_novo.fasta.

To identify contigs associated with different genomic compartments, three genomes of *Francisella* Endosymbiont (FLE) associated with ticks (F-Om (GCA_003069505.1), FLE-Om (GCA_002095075.2), FLE-Am (GCA_001753795.1)] and five *Hyalomma* mitochondrial genomes (NC_056189, MW546283, KY457529, MF101817, NC_062166) available in public databases were used to build reference datasets. These were employed in blastn searches to classify assembled contigs according to their likely origin. For nuclear and *Francisella* assemblies, genome completeness was assessed with BUSCO v5.8.2 (Manni et al. 2021) using the acari_odb12 and gammaproteobacteria_odb12 datasets, respectively (Tegenfeldt et al., 2025). Assembly statistics and quality metrics were assessed and visualized using BlobToolskit v2.5.0 (Challis et al., 2020). Genomic features, including scaffold count and length, N50, GC content, and sequence gaps, were summarized in a snail plot. This visualization also incorporates the BUSCO assessment to provide a comprehensive overview of the assembly’s completeness and structural integrity.

### Genome annotation

Functional genome annotation was performed with the external NCBI Eukaryotic Genome Annotation Pipeline (EGAPx) v0.4.1-alpha (Thibaud-Nissen et al. 2016) (https://github.com/ncbi/egapx). The appropriate NCBI taxonomic identifier (taxid 34625, corresponding to the genus *Hyalomma*) was provided, together with high-quality RNA-seq data from the same individual whole organism to support transcript evidence (BioSample SAMEA114541064; Experiment ERX13630423).

Gene-space completeness was assessed for the *Hyalomma lusitanicum* genome and the 11 chromosome-level genome assemblies available for Ixodida (NCBI, genome database: MW546280, MW546283, MF101817, OM368315, OM368316, MW546284, MT270687, MW366628, NC_062166, OM368314, KY457529) using BUSCO proteome with the acari_odb12 dataset. In addition, ortholog-based quality assessment was performed using OMArk (Nevers et al. 2024) with the Metazoa dataset. OMArk automatically selected the Ixodidae lineage to evaluate Hierarchical Orthologous Groups (HOGs) in a lineage-specific context.

Mitochondrial genome was performed using MITOS (Bernt et al., 2013). The new annotation was compared with the available *Hyalomma* mitogenome to corroborate the protein-coding frame and the completeness of tRNAs.

### Annotation of the Francisella genome

Genome annotation was performed with the NCBI Prokaryotic Genome Annotation Pipeline (PGAP) 2025-05-06.build7983 (Tatusova et al. 2016; Haft et al. 2018) using default parameters, specifying *Francisella* as the target genus. Intact coding genes and pseudogenes were identified using the ‘Annotate’ command in Pseudofinder (Syberg-Olsen et al. 2022).

### Tick synteny

To evaluate chromosomal conservation and structural evolution, a pairwise synteny analysis was performed between the *de novo* assembly and the reference genome of *Rhipicephalus microplus* (GCF_043290135.1). Orthologous protein clusters were identified by estimating pairwise similarity among proteomes using Diamond BlastP (Buchfink et al. 2021). The resulting alignments were used as input for MCScanX v1.0.0 (Wang et al., 2012) to identify collinear blocks requiring at least five genes per block. Syntenic relationships were visualized using the SynVisio web server (https://synvisio.github.io/#/) (Bandi and Gutwin, 2020). For visualization, *R. microplus* was set as the reference, comprising 10 autosomal chromosomes (rm1–10) and the X sexual chromosome (rmX), while our assembly was represented by the 18 largest contigs (hl1–18). In the resulting dual synteny plot, blue ribbons indicate conserved gene orientation (collinear), whereas red ribbons represent inverted orientations between the two genomes.

### Phylogenetic trees

Evolutionary trees for *Hyalomma lusitanicum* and *Francisella* endosymbiont were inferred using Maximum-Likelihood (ML) under the best-fit evolutionary mixture models. To that, single-copy genes at amino acid codification were inferred using BUSCO for both genomes using acari_odb12 and gammaproteobacteria_odb12, respectively. Then, we perform the orthology search with Orthofinder (Emms and Kelly, 2019). The selected homologous genes were first produced with PREQUAL (Whelan et al., 2018) to mask residues of the genes without evidence of homologous position to be aligned and filtered using MAFFT (Katoh and Standley, 2013) and BMGE (Criscuolo and Gribaldo, 2010). Final matrices were used to infer the evolutionary trees with IQ-TREE (Minh et al, 2020) under empirical mixture models of Modelfinder (Kalyaanamoorthy et al., 2017). Best-fit evolutionary mixture models were selected using *-m TESTONLYNEW -madd C10,C20,C30,C40,C50,C60* under BIC (Schwarz, 1978). *Neoseiulus* and *Phytoseiulus* (Mesostigmata) species were used as outgroups based on recent phylogenetic hypotheses (Bhoi et al., 2026).

## RESULTS AND DISCUSSION

Within the framework of the ERGA–BGE initiative, we generated and annotated high-quality nuclear and mitochondrial reference genomes for *Hyalomma lusitanicum*, a species with potential relevance in the circulation of Crimean–Congo hemorrhagic fever virus (CCHFV) in the Iberian Peninsula. In addition, we assembled and annotated the complete genome of its associated *Francisella* Endosymbiont (FE), further expanding the resources available for investigating tick-symbiont interactions.

Genomic resources for *Hyalomma lusitanicum* and ticks in general remain limited, with only 11 genomes assembled at the chromosome level (Figure 1). Here, we present a high-quality genome assembly that substantially improves the resolution for investigating tick genome structure. Although chromosome-level scaffolding was not achieved, our nuclear assembly is among the most contiguous currently available, representing a highly complete genome resolved into a limited number of scaffolds.

**Figure 1.**
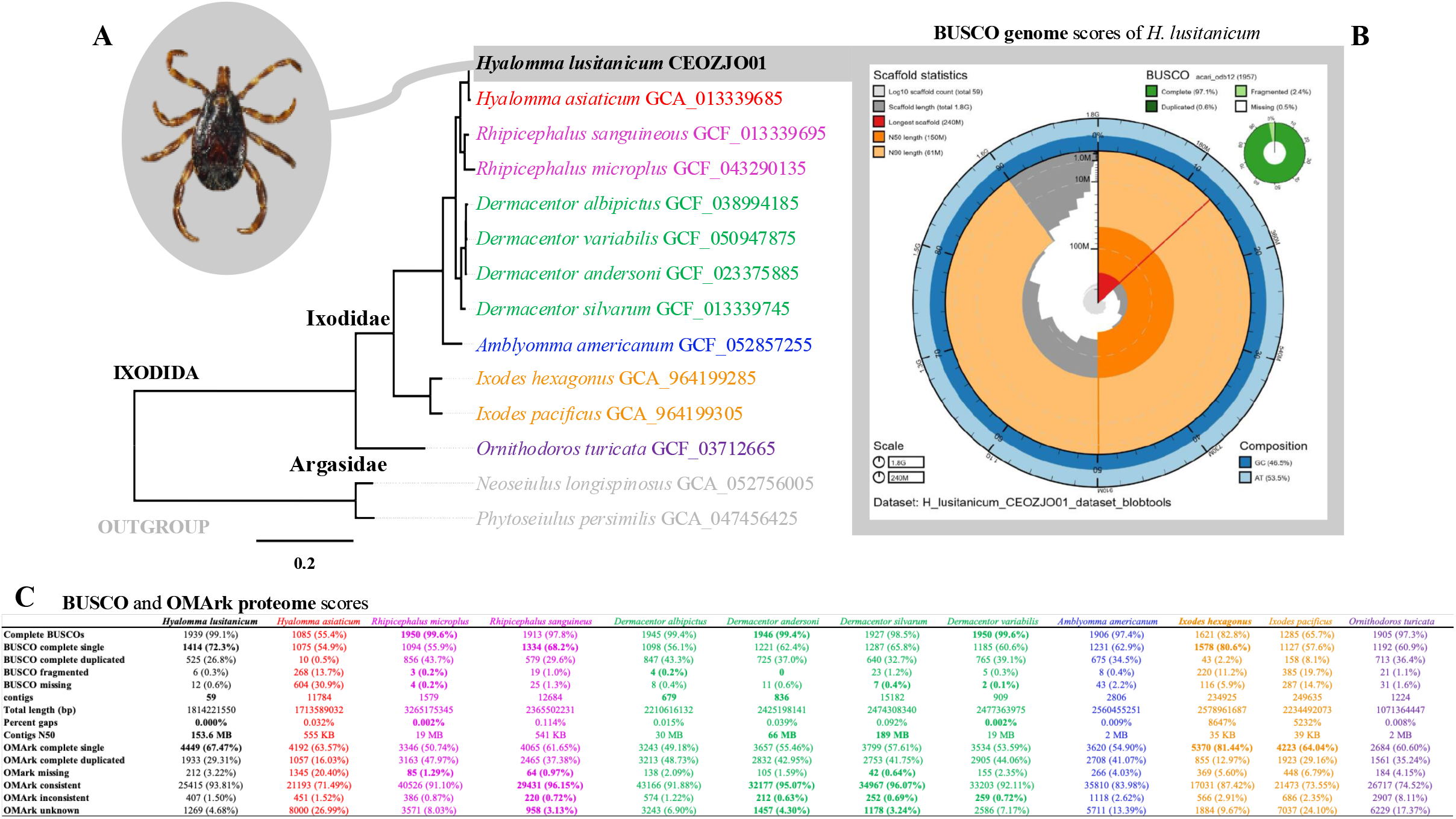
**A.** Maximum likelihood phylogenomic tree of Ixodida, including all available chromosome-level genomes from NCBI. All nodes were fully supported. A photograph of a *Hyalomma lusitanicum* individual collected near the genome sampling locality is shown alongside the tree. **B**. Snail plot summarizing assembly statistics, including scaffold metrics, contig size distribution, BUSCO completeness of the genome based on the acari_odb12 database, and overall genome composition. **C**. Comparative table of annotated proteome completeness, also using the acari_odb12 database, across chromosome-level genomes. The three highest scores per metric are highlighted in bold.

Such high-quality assemblies provide a robust framework to explore gene family evolution, detect signatures of natural selection, and infer demographic history, all of which are essential for understanding the evolutionary dynamics of the species (Theissinger et al., 2023). In a One Health context, integrating genomic, ecological, and epidemiological data facilitates the identification of candidate genes associated with acaricide resistance, host-parasite interactions, and environmental adaptation (Tigistu-Sahle et al., 2023), ultimately supporting the development of more effective, evidence-based strategies for vector surveillance, management, and control.

### Hyalomma lusitanicum genome assembly

The final genome assembly of *Hyalomma lusitanicum* spans a total length of 1.81 Gb (1,814,221,550 bp) and is highly contiguous, comprising only 59 scaffolds (with 111x and 98x of depth coverage to short and long reads, respectively). The assembly exhibits a GC content of 46.53%, consistent with other ixodid tick genomes (43.7%, SD 4.9; https://www.ncbi.nlm.nih.gov/datasets/genome/?taxon=6935). Contiguity metrics further support its high quality, with a scaffold N50 of 153.6 Mb and an L50 of 5, indicating that half of the genome is contained within just five scaffolds. Notably, the assembly is gap-free, with zero unresolved regions, reflecting its completeness and structural continuity (Figure 1). Contig-level statistics (N50 and L50) also indicate a high degree of base-level contiguity. Genome completeness, as assessed by BUSCO v5.8.2 (Manni et al. 2021) with the acari_odb12 dataset, yielded 97.1%, suggesting a high recovery of conserved single-copy orthologs (Figure 1B). In addition, k-mer-based evaluation showed that 78.24% of k-mers are represented in the assembly, with a consensus quality value (QV) of 65.28 estimated with Merqury (Rhie et al., 2020), indicating high base accuracy. Together, these metrics demonstrate that this assembly represents a near-complete, high-quality genomic resource at the chromosome scale.

However, our assembly resulted in 59 contigs, of which 18 were identified as putative chromosomes, exceeding the 11 chromosomes inferred as ancestral by GoaT. This discrepancy is likely driven by incomplete Hi-C data, which limited the accurate scaffolding of contigs into full chromosomal sequences. Synteny analyses against the most complete *Rhipicephalus microplus* genome available further support this interpretation, as multiple *H. lusitanicum* scaffolds map to single ancestral chromosomes (*e*.*g*., hl1, hl3, and hl12 align to the rmX sex chromosome). Together, these results indicate that several contigs likely represent fragmented portions of the same chromosome (Figure 2).

**Figure 2.**
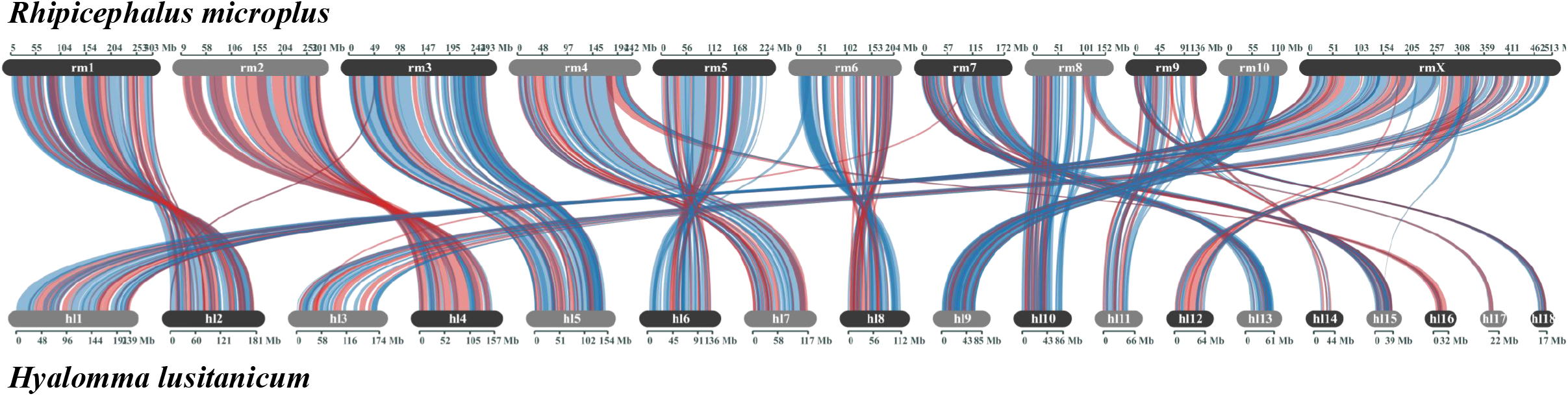
Dual synteny plot between *Hyalomma lusitanicum* and *Rhipicephalus microplus* reference genome. The syntenic relationships were identified based on orthologous protein clusters using MCScanX. The reference genome of *R. microplus* (top) is represented by its 10 autosomal chromosomes (rm1-10) and the X sexual chromosome (rmX). The *de novo* assembly of *H. lusitanicum* (bottom) is represented by the 18 largest scaffolds (hl1-18). Each ribbon represents a collinear block containing at least five genes. Blue ribbons indicate conserved gene orientation (collinearity), while red ribbons highlight genomic inversions. The scale on the chromosomes and scaffolds is indicated in megabases (Mb).

The mitochondrial genome of *Hyalomma luitanicum* was isolated from the SAMEA114541046 biosample (NCBI number). The circular molecule is 14,752 bp in length and contains 37 genes: 22 transfer RNAs (tRNAs), 13 protein-coding genes (CDSs), and two ribosomal RNA genes (rRNAs) (NCBI accession number). As observed in most hard-tick mitochondrial genomes, it includes two putative control regions (Uribe et al., 2020). The gene organization of the mitochondrial genome is shown in Figure 1S.

### Hyalomma lusitanicum genome annotation

The genome annotation identified a total of 20,638 protein-coding genes, giving rise to 35,134 transcripts. In addition, 1,422 non-coding genes were annotated, including rRNA, tRNA, lncRNA, snoRNA, and snRNA elements. A total of 5,775 pseudogenes were also detected, reflecting the extent of gene duplication and degeneration processes within the genome. Annotation quality metrics based on BUSCO and OMArk proteome analyses indicate high completeness, comparable to the highest-quality tick genomes currently available, including those of *Dermacentor, Rhipicephalus, Amblyomma*, and *Ornithodoros* (Figure 1C).

### Ticks phylogenomic tree

Protein genes at the amino acid level were used to reconstruct the evolutionary tree under the Q.insect+F+I+R4 evolutionary model (*best log-likelihood* −2246433.921). Our phylogenomic hypothesis recovered *Hyalomma* as monophyletic and sister to *Rhipicephalus*, as previous phylogenetic trees (Charrier et al., 2019). *Ixodes* is recovered as the first divergence of Ixodidae (hard-ticks), which was the sister of Argasidae (soft ticks), that was represented by *Ornithodoros* (Figure 1A). Future studies have to be addressed, including denser genomic taxon sampling to evaluate the deeper phylogenetic relationships of Ixodida.

### Francisella Endosymbiont (FE) genome assembly, annotation, and phylogeny

From the ERR13381541 sequencing data, we assembled the first complete *Francisella* endosymbiont (FE) associated with ticks. The circular genome is 1,511,359 bp in length, with a mean coverage of 180× (SD = 23.6), and comprises 1,679 genes. Of these, 731 were identified as pseudogenes, while 10 correspond to ribosomal RNA (rRNA), 37 to transfer RNA (tRNA), and four to other non-coding RNAs (ncRNAs) genes (Figure 3B). The high number of pseudogenes, together with the reduced genome size compared to non-endosymbiotic *Francisella* species (Figure 3), is consistent with patterns widely reported in obligate endosymbionts and is likely driven by long-term host dependence and genome reduction (Nardi et al., 2021; Echeverry-Pérez et al., 2025).

**Figure 3.**
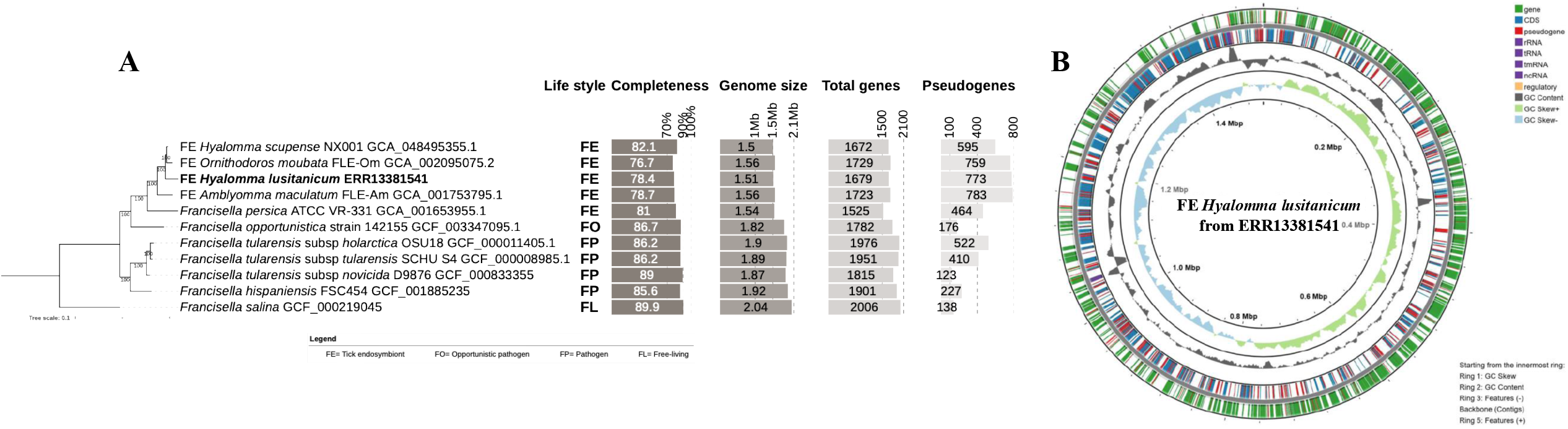
**A.** Maximum likelihood phylogenomic tree of the *Francisella* genus, including the most representative publicly available genomes of different lifestyles in NCBI. The tree was inferred with 174 protein-coding genes (58,868 amino acid positions) using the best-fit evolutionary model (Q.plant+F+I+R2). All nodes were fully supported. Comparative scores of the lifestyle, completeness, genome size, total genes, and pseudogenes are shown alongside the tree. **B**. Circular genomic map of the FE from *Hyalomma lusitanicum* assembly. The visualization was constructed using Proksee (Grant et al. 2023) based on PGAP and Pseudofinder-derived annotations. Concentric rings from the innermost to the outermost track represent: (1) positive (light green) and negative (light blue) GC skew; (2) GC content (gray); (3) negative and (4) positive strands including features such as RNAs (purple), regulatory genes (peach), pseudogenes (red), CDS (blue), and genes (green). The scale of the chromosome is indicated in megabases (Mb).

The FE strains were recovered as monophyletic with *F. persica* as the first divergence, followed by the FE strain of *Amblyomma maculatum*. FE strains *Hyalomma* are different lineages, with FE of *H. scupense* sister clade of *Ornithodoros moubata* and FE strain of *Hyalomma lusitanicum* sister lineage of them (Figure 3A).

## CONCLUSIONS

We present one of the most contiguous and compact genomic resources currently available for ticks, both in terms of scaffold-level assembly and gene annotation. The high contiguity of the nuclear genome, together with the complete mitochondrial and endosymbiont genomes (the first complete genome for *Francisellas* endosymbiont bacteria associated to ticks), provides a comprehensive genomic framework for *Hyalomma lusitanicum*. Given the ecological relevance of this species in the western Mediterranean and its potential role in the transmission of pathogens such as CCHFV, these resources will be instrumental for investigating patterns of local adaptation, genomic structure, and population dynamics across its distribution, as well as for identifying genes under selection that may influence vector competence. Beyond their evolutionary and genomic relevance, these resources will directly support future studies aimed at developing surveillance, control, and management programs for this species of growing public health concern.

## Supporting information

NONE

## CRediT AUTHORSHIP CONTRIBUTION STATEMENT

NE, RF, AsB, and RM provided support in sampling, shipping of biological material, metadata collection, and management; LA and MG extracted DNA, prepared libraries, and performed sequencing; FCF, FC, and JG-G performed genome assembly and curation under the supervision of TSA; CdG helped generate the report. JSEP and JEU performed the formal analyses. JEU wrote the initial draft, and all authors checked and agreed with it.

## DECLARATION OF COMPETING INTEREST

None.

## DATA AVAILABILITY

*Hyalomma lusitanicum* and the related genomic study were assigned to Tree of Life ID (ToLID) 49205, and all sample, sequence, and assembly information are available under the umbrella BioProject PRJEB77820. The sample information is available at the following BioSample accessions at NCBI: SAMEA114541046, SAMEA114541064, and SAMEA114541056. The experiment accession numbers are the following: ERX14823192, ERX13630423, ERX13167078, and ERX12752434, and the genome assembly is accessible from ENA under the accession number GCA_981163835. The annotated genome is available at the Ensembl website (https://projects.ensembl.org/erga-bge/). Documentation related to the genome assembly and curation can be found in the ERGA Assembly Report (EAR) document available at (https://github.com/ERGA-consortium/EARs/blob/main/Assembly_Reports/Hyalomma_lusitanicum/HyaLusi11/qqHyaLusi11_EAR.pdf). Further details and data about the project are hosted on the ERGA portal at https://portal.erga-biodiversity.eu/data_portal/49205.

## ACKNOWLEDGMENTS

Thanks to Drago (https://aic.csic.es/supercomputador-drago/) and CESGA (https://www.cesga.es/infraestructuras/computacion/) supercomputers, and the Smithsonian Institution High Performance Computing Cluster (https://doi.org/10.25572/SIHPC), where part of the high-performance computing was conducted.

## SUPPLEMENTARY MATERIALS

Figure 1S. Mitochondrial phylogeny

---

**Figure 1S.** Mitochondrial phylogeny based on a matrix with 12587 nucleotide positions, ML inference, and the best-fit evolutionary model (GTR+F+I+R3; *best log-likelihood* −84713.021). *Hyalomma lusitanicum* is strongly recovered as closely related to *H. truncatum* and *H. anatolicum*, but the relationships between them are weakly supported. Previous analyses have also recovered these close relationships (Sands et al., 2017). However, this poor resolution could be due to a scarce taxon sampling of a fast radiation in the isolation of those related species.

## Notes

### Competing Interest Statement

The authors have declared no competing interest.

