## Supplementary figures and images for "ERGA-BGE reference genomes of *Hyalomma lusitanicum* and its obligate *Francisella* endosymbiont as a genomic resource for One Health research"

### NONE

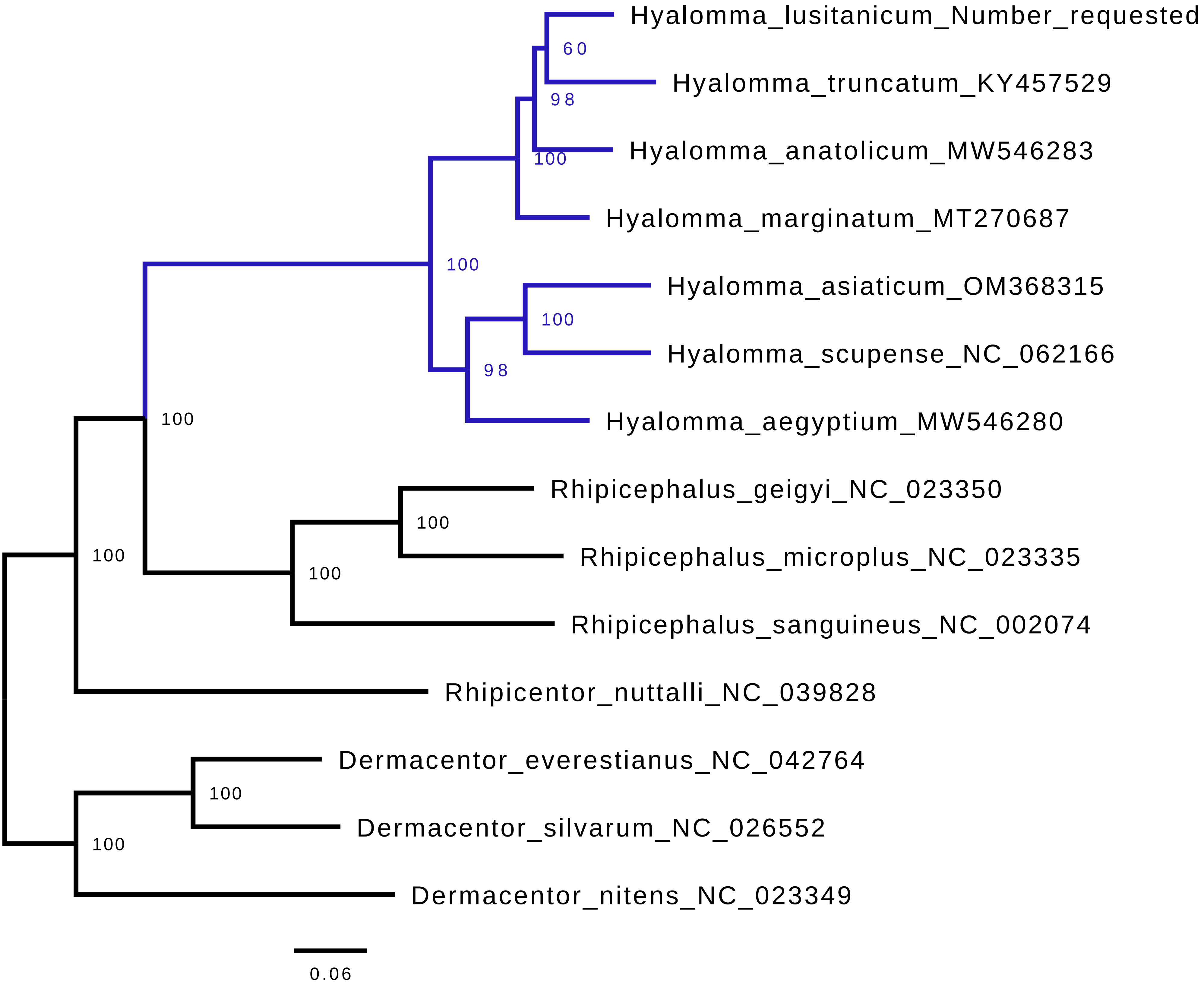
